# Chemical genetic interactions elucidate pathways controlling tuberculosis antibiotic efficacy during infection

**DOI:** 10.1101/2024.09.04.609063

**Authors:** Peter O. Oluoch, Eun-Ik Koh, Megan K. Proulx, Charlotte J. Reames, Kadamba G. Papavinasasundaram, Kenan C. Murphy, Matthew D. Zimmerman, Véronique Dartois, Christopher M. Sassetti

## Abstract

Successful tuberculosis therapy requires treatment with an unwieldy multidrug combination for several months. Thus, there is a growing need to identify novel genetic vulnerabilities that can be leveraged to develop new, more effective antitubercular drugs. Consequently, recent efforts to optimize TB therapy have exploited Mtb chemical genetics to identify pathways influencing antibiotic efficacy, novel mechanisms of antibiotic action, and new targets for TB drug discovery. However, the influence of the complex host environment on these interactions remains largely unknown, leaving the therapeutic potential of the identified targets unclear. In this study, we leveraged a library of conditional mutants targeting 467 essential Mtb genes to characterize the chemical-genetic interactions (CGIs) with TB drugs directly in the mouse infection model. We found that these *in vivo* CGIs differ significantly from those identified *in vitro*. Both drug-specific and drug-agnostic effects were identified, and many were preserved during treatment with a multidrug combination, suggesting numerous strategies for enhancing therapy. This work also elucidated the complex effects of pyrazinamide (PZA), a drug that relies on aspects of the infection environment for efficacy. Specifically, our work supports the importance of coenzyme A synthesis inhibition during infection, as well as the antagonistic effect of iron limitation on PZA activity. In addition, we found that inhibition of thiamine and purine synthesis increases PZA efficacy, suggesting novel therapeutically exploitable metabolic dependencies. Our findings present a map of the unique *in vivo* CGIs, characterizing the mechanism of PZA activity *in vivo* and identifying novel targets for TB drug development.

**Significance:** The inevitable rise of multi-drug-resistant tuberculosis underscores the urgent need for new TB drugs and novel drug targets while prioritizing synergistic drug combinations. Chemical-genetic interaction (CGI) studies have delineated bacterial pathways influencing antibiotic efficacy and uncovered druggable pathways that synergize with TB drugs. However, most studies are conducted *in vitro*, limiting our understanding of how the host environment influences drug-mutant interactions. Using an inducible mutant library targeting essential Mtb genes to characterize CGIs during infection, this study reveals that CGIs are both drug-specific and drug-agnostic and differ significantly from those observed *in vitro*. Synergistic CGIs comprised distinct metabolic pathways mediating antibiotic efficacy, revealing novel drug mechanisms of action, and defining potential drug targets that would synergize with frontline antitubercular drugs.

## Introduction

Tuberculosis (TB), caused by the bacterial pathogen *Mycobacterium tuberculosis* (Mtb), remains a leading cause of global mortality attributable to a single infectious disease, with 1.3 million deaths reported in 2022 (1). Despite the general success of the ‘short-course’ combination therapy (2), TB treatment remains long and arduous and commonly results in unfavorable outcomes (3). Thus, there is an urgent need to develop effective, short, and well-tolerated antitubercular regimens. The recent success of a novel oral regimen for drug-resistant TB (4, 5) has provided momentum for expanding the anti-TB drug pipeline, focusing on repurposing, repositioning, or identifying first-in-class drug candidates (6). Furthermore, the seemingly inevitable emergence of resistance, including to the recently introduced anti-tubercular drugs (7–9), underscores the need to continuously replenish the TB drug pipeline and identify novel drug targets.

Understanding and predicting how an antibacterial drug will perform at the site of infection represents a major challenge for development efforts. The host environment has been shown to profoundly affect bacterial metabolism and antibiotic activity (10–12). This is evident in the activity of single drugs, as both immune pressure and the nutritional environment experienced during infection can reduce the potency and killing rates of many drugs (11, 13). Conversely, the activity of pyrazinamide (PZA), a cornerstone component of TB therapy credited with shortening treatment duration (14, 15), is accentuated in the infection environment. This drug inhibits the coenzyme A (CoA) synthesis pathway essential for bacterial growth both *in vitro* and during infection (16–18). However, PZA has very poor activity in standard *in vitro* culture conditions, and its potency depends on the low pH and possibly other conditions encountered during infection. This remarkable difference in potency has left the mechanism(s) underlying PZA’s clinical effectiveness unclear. These examples further highlight the difficulty in using the *in vitro* data produced during drug development to predict clinical potency.

Given the profound effects of changing environmental conditions on the activity of single drugs, it is not surprising that this also impacts their interactions in combination (19). Historically, successful tuberculosis treatment has relied on a combination of multiple drugs (20, 21), and current efforts to identify new regimens focus on synergistic combinations that could increase drug potency and improve treatment outcomes (22–24). Both experimental and computational tools have been used to create comprehensive catalogs of drug-drug interactions (25–27). Complementary genetic approaches including transposon insertion sequencing (Tn-seq), CRISPR interference, and conditional proteolysis have been developed to substitute drugs with mutants (28–30), enabling more comprehensive chemical-genetic interaction (CGI) studies. Each of these approaches indicates that interactions between inhibitors are very sensitive to environmental conditions. For example, similar Tn-seq methods were used to identify interactions between non-essential Mtb genes and drugs both *in vitro* (31) and in infected mice (32) and detected very different sets of CGIs in the two conditions. CGI and drug-drug interaction studies using different defined media conditions could attribute some of these differences to carbon source utilization or pH, but no simple *in vitro* culture condition adequately models the complex infection environment (13, 23). Thus, methods to interrogate drug mechanisms of action and the potential synergistic activities between drugs directly in the infection environment are needed to focus drug development efforts on the most promising targets and mechanisms.

In this work, we sought to identify CGIs relevant to the infection environment using a library of conditional Mtb knockdown mutants targeting genes that are essential for Mtb growth. This set of genes contains the most promising potential new drug targets. However, essential genes cannot be assessed via Tn-seq and therefore have been excluded from previous studies (32). We characterized interactions between these essential functions and components of the first-line TB drug regimen directly in a mouse model of infection and treatment. This work shows that *in vivo* CGIs are distinct from those identified *in vitro* and are mostly drug-specific. However, we also identified a small number of drug-agnostic synergies and found that even drug-specific interactions are often evident during multi-drug treatment. We also report on novel PZA-specific synergies, supporting the proposed mechanism of action for this drug and characterizing a new link between PZA and thiamine and purine metabolism. These findings underscore the involvement of multiple pathways and mechanisms controlling antibiotic efficacy and identify attractive targets for TB drug discovery.

## Results

### A genetic approach to define chemical-genetic interactions during infection

To characterize chemical-genetic interaction signatures during infection, we leveraged an inducible hypomorph system designed to conditionally target essential proteins for proteolytic degradation (13, 29, 33). A DAS+4 tag (DAS-tag) was engineered at the 3’end of 467 essential genes, targeting the proteins for anhydrotetracycline (ATc)-inducible Clp protease degradation (29). This represents ∼75% of genes previously shown to be essential for *in vitro* Mtb growth (18, 34). For each tagged gene, four hypomorphs were generated to express different levels of the Clp adapter protein, SspB, and produce distinct levels of protein knockdown. SspB versions 2, 6, 10, and 18 represent increasing strengths of knockdown. Each mutant was engineered to include a unique 20-nucleotide barcode sequence, allowing the quantification of strain abundance in a mixed pool (**Fig. 1A**).

**Figure 1:**
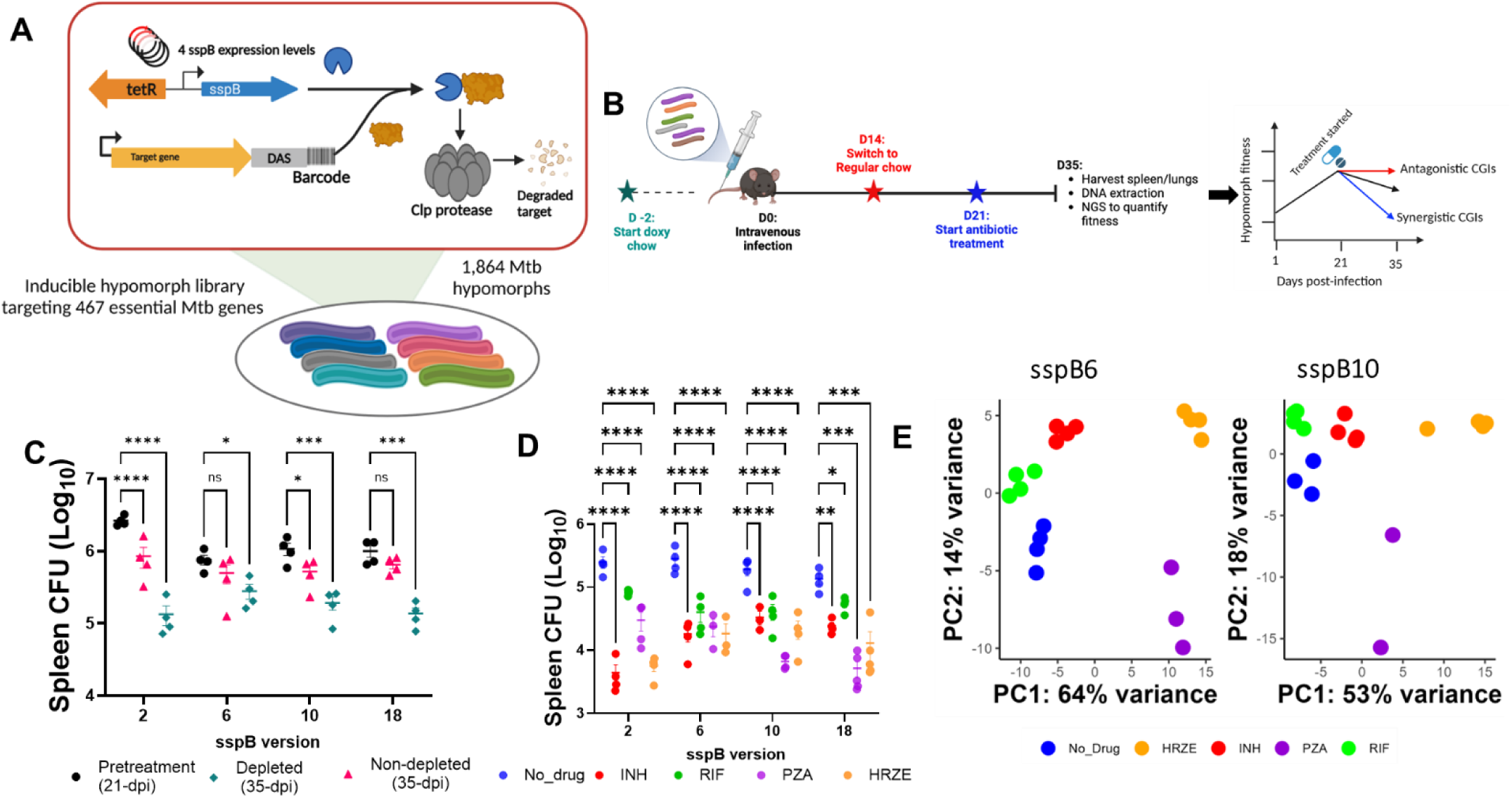
Genetic strategy for *in vivo* chemical-genetic interaction profiling in M. tuberculosis. **(A)** Design of the conditional hypomorph library as previously described <u>(22)</u>, targeting 467 essential Mtb genes at four sspB levels of protein depletion and a total of 1864 hypomorph strains**. (B)** Illustration of the *in vivo* CGI screen (See methods). **(C)** Comparison of spleen CFU in non-treatment control groups across all the sspB levels (n = 4 biological replicates, mean ± s.e.m). **(D)** Comparison of spleen CFU in depleted-treated vs depleted-untreated mice groups across all sspB depletion levels (n = 2-4 biological replicates, mean ± s.e.m). Results from two-way ANOVA with Dunnett’s multiple comparisons test; *P < 0.05, **P < 0.01, ***P < 0.001, ****P < 0.0001. **(E)** Principal component analysis (PCA) scores plot of normalized mutant barcode counts in depleted-treated and non-treated mice groups at sspB #6 and #10.

We designed an experimental strategy that integrated mouse infection, control of hypomorph depletion, optimized drug dosing, and quantification of bacterial killing during antibiotic treatment to elucidate chemical-genetic interactions with first-line antitubercular drugs (**Fig. 1B**). Mice were infected via the intravenous route (IV) with hypomorph pools (∼100 CFUs/strain) corresponding to each of the sspB levels. All mice were fed a doxycycline-containing diet for two weeks to allow the establishment of infection in the absence of gene knockdown, then transitioned to a regular diet to initiate specific protein depletion for an additional week before antibiotic treatment. Non-depleted control mice were maintained on the doxycycline-containing diet for the entire study period. At 21 days postinfection (21-dpi), groups of animals were treated with isoniazid (INH), rifampicin (RIF), pyrazinamide (PZA), or combination therapy consisting of these three drugs plus ethambutol (HRZE) for two weeks (**Fig. 1B**). Surviving bacteria were then recovered from the spleen, and the relative abundance of each mutant was quantified via barcode amplification and sequencing. Based on previous studies, the spleen represents a robust model of Mtb intracellular growth in the presence of a strong adaptive immune response (17, 32, 34–36), where the establishment of large mutant libraries is possible.

In this study, antibiotics were administered via drinking water. To ensure that the antibiotic concentrations were relevant to human dosing during infection and remained stable during administration, we quantified the concentration of all four antibiotics alone or in the HRZE combination in drinking water (**Fig. S1A-B**) and monitored the plasma concentration of each drug in healthy mice (See Methods). Using these data, we were able to achieve plasma drug exposure (24-hour area under the curve) (**Fig. S1C**) comparable to the ranges observed in adult humans (37). To validate our model system, we independently quantified the effects of protein depletion and drug treatment on the bacterial load. The efficacy of *in vivo* depletion of targeted proteins was assessed by comparing bacterial clearance across all depletion levels in the absence of treatment. As expected with the inhibition of genes essential for *Mtb* growth, hypomorph depletion resulted in a significant decrease in bacterial burden in the spleen between 21- and 35-days post-infection (dpi) (**Fig. 1C**). Similarly, each antibiotic treatment independently resulted in a significant reduction in bacterial burden in the spleen across all depletion levels (**Fig. 1D**). These results confirmed that the effects of protein depletion and drug treatment can be combinatorically assessed.

To define the effects of each drug on individual mutants in the hypomorph library, we recovered surviving bacteria in the spleen by plating ∼10^6 colony-forming units (CFU) followed by barcode amplification, deep sequencing, and quantification. Overall, the distribution of hypomorph barcode counts across all *in vivo* conditions was comparable (**Fig. S2A**) and significantly correlated across biological replicates (Pearson correlation r > 0.87, *P* < 0.001; **Fig. S2B**). Raw barcode counts were filtered (Methods) to exclude low-abundance hypomorphs lost from the pool due to infection bottlenecks (38), retaining 1140 out of the 1864 Mtb hypomorphs (317 sspB-2; 319 sspB-6; 282 sspB-10; 222 sspB-18). Principal component analysis (PCA) was applied to the filtered dataset to examine the effect of each drug treatment on the composition of the hypomorph library pools selected in individual mice. In this PCA space, mice treated with the same antibiotic clustered together (**Fig. 1E**). The first principal component (PC1) accounted for 53-64% of the variance in the library, aligning the effect of PZA alone or in the HRZE combination. PC2 accounted for 14-18% of the variance and delineated the effects of both INH and RIF alone, as well as in the HRZE combination (**Fig. 1E**). Taken together, we conclude that both the inducible knockdown system and the antibiotic treatment worked independently during infection, which should allow the combined effects of these perturbations (chemical-genetic interactions) to be assessed.

### CGIs are both drug-specific and drug-agnostic

To characterize the differential effects of antibiotics on individual mutants, we implemented a DEseq-based differential barcode representation analysis (DEBRA) (39), comparing barcode counts in DAS-tag depleted and drug-treated groups vs those that were depleted in the absence of drug therapy. In this analysis, only CGIs with effects that significantly exceed additivity are detected. The resulting interactions summarized as log_2_ fold changes (LFC), and FDR-adjusted *P* values across all conditions are shown in **Fig. S2C** and listed in **Supplementary Table 1**. Overall, we observed both treatment- and depletion-level-dependent differences in the frequency of CGI (**Fig. 2A**). “Synergistic” and “antagonistic” interactions were defined as gene depletions that significantly increased or decreased drug efficacy, respectively. To determine whether the CGIs across different treatment groups were influenced by the replication/growth rates of the hypomorphs, we assessed the relationship between mutant abundance in each treatment (Log_2_FC, treated vs. non-treated) compared to growth controls (Log_2_FC, depleted vs. non-depleted). We observed no relationship between growth rate and synergistic or antagonistic CGIs across all treatment groups (**Fig. 2B**), indicating that the observed CGIs are not primarily dependent on the replication rates of individual mutants, and appeared to reflect specific interactions between gene knockdown and drug treatment.

**Figure 2:**
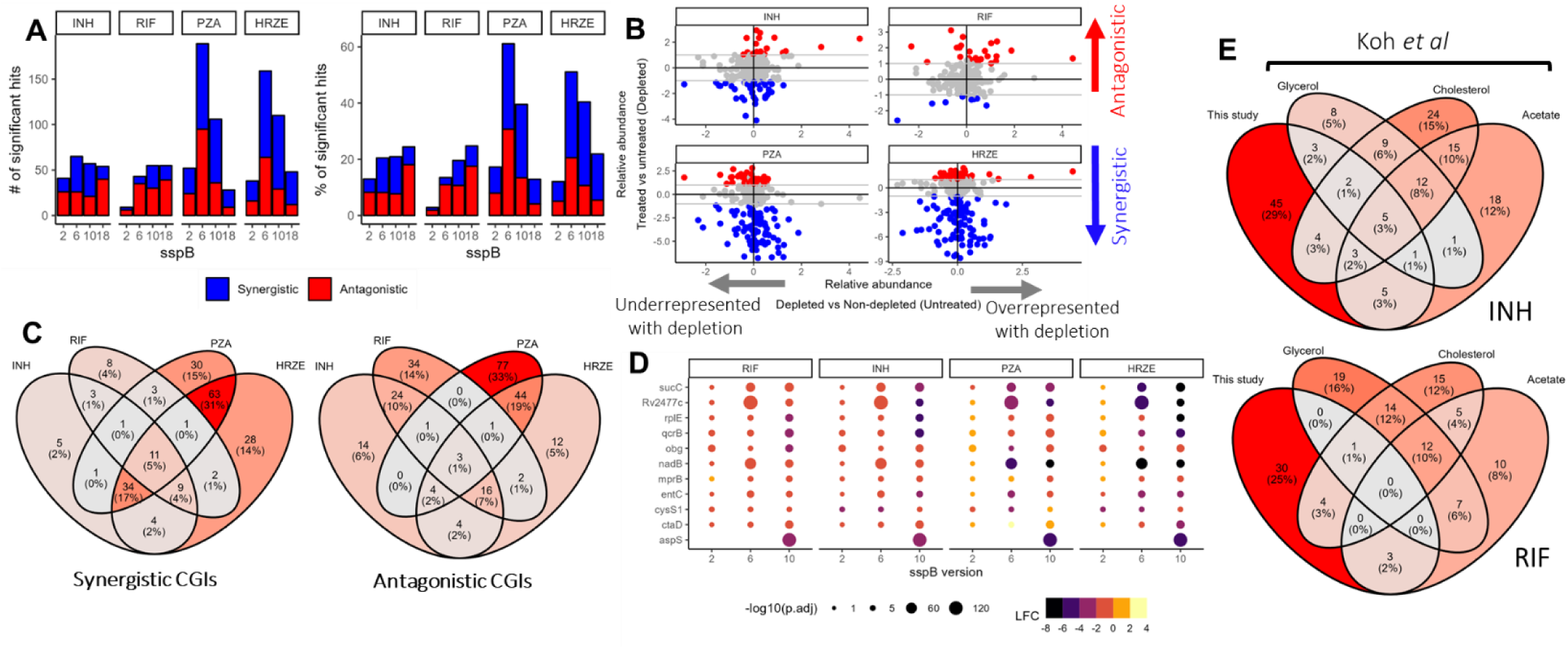
Antibiotic-specific and antibiotic-agnostic CGI signatures during infection. **(A)** Number and percentage of significant synergistic and antagonistic interactions across different treatment conditions. **(B)** The relative abundance, expressed as log2 fold change, for different hypomorph strains in untreated controls (x-axis) and treated groups (y-axis). Mutants with synergistic CGIs are in blue, while antagonistic CGIs are in red. **(C)** Venn diagram of the relationship of genes with synergistic or antagonistic CGIs in at least 1 depletion level**. (D)** Bubble plots of log2 FC for mutants with synergistic CGIs in all treatment conditions. Plot size is scaled by the log_10_(FDR) values while the color range represents the log_2_ FC values. **(E)** Venn diagram of the relationship CGIs during *in vivo* infection vs *in vitro* culture conditions with three different carbon sources.

Next, we analyzed the dataset to define interaction patterns across different conditions. Most synergistic interactions across all depletion levels occurred under PZA (144/1084) and HRZE (152/1107) conditions, which also exhibited the largest overlap in unique synergistic and antagonistic CGIs. Overall, 44% and 34% of the synergistic and antagonistic PZA interactions, respectively, were also evident in the HRZE combination (**Fig. 2C**). On the other hand, isoniazid and rifampicin conditions had the lowest number of synergistic CGIs (68/1113 and 38/1125, respectively). As with PZA, some of these synergistic interactions, 13 out of 68 in isoniazid and 11 out of 38 in rifampicin, overlapped with synergistic HRZE CGIs (**Fig. 2C; Supplementary Table 2**). Thus, while most CGIs found for individual drugs were specific to that condition, many of these were also apparent in combination therapy, suggesting that they could be leveraged to enhance the effects of a multidrug regimen.

In addition to the predominant drug-specific effects, we also observed mutants with broadly synergizing effects across multiple treatment conditions. Thirty-four mutants resulted in synergistic CGIs in INH, PZA, and HRZE, while an additional nine mutants synergistically interacted with INH, RIF, and HRZE (**Fig. 2C; Supplementary Table 2**). We also observed eleven mutants with drug-promiscuous synergies, with knockdowns in these genes resulting in synergistic interactions across all treatment conditions. This set of drug-agnostic mutants included genes in multiple pathways, including respiration (*qcrB* and *ctaD*), protein synthesis (*aspS*, *cysS1*, Rv2477c (*ettA*) and *obg*), and NAD biosynthesis (*nadB*) (**Fig. 2D**). Thus, while the majority of CGIs are drug-specific, a small subset shows drug-agnostic CGI profiles and likely act by altering fundamental aspects of bacterial physiology.

We next sought to determine if the observed interactions were also apparent under *in vitro* culture conditions or if they were unique to the *in vivo* environment. The *in vivo* dataset was compared to a previous *in vitro* CGI screen conducted using the same hypomorph library that was exposed to MIC-level concentrations of antibiotics in three different media compositions with glycerol, acetate, or cholesterol as the major carbon sources (13). We observed considerable overlap in synergistic CGIs in both INH (23 hypomorphs shared between 70 *in vivo* and 110 *in vitro*) and RIF (8 hypomorphs shared between 40 *in vivo* and 90 *in vitro,* **Fig. 2E**). However, these overlaps did not reach statistical significance (INH: *P* = 0.4624, RIF: *P* = 0.8416 by Fisher’s Exact Test). We also observed a similar degree of overlap in antagonistic CGIs in INH (18 hypomorphs shared out of 61 *in vivo* and 131 *in vitro, P* = 0.158, Fisher’s Exact Test), and RIF (34 hypomorphs shared out of 81 *in vivo* and 170 *in vitro, P* = 0.616, Fisher’s Exact Test, **Fig. S2D**). Overall, neither the union of *in vitro* data nor any single *in vitro* condition served as a significant predictor of *in vivo* CGI. Thus, most CGIs are unique to the *in vivo* or *in vitro* environments, and the simple *in vitro* culture conditions investigated in these datasets are not predictive of bacterial killing rate during infection.

### CGIs are reproducible during pulmonary infection

While splenic tuberculosis serves as a suitable model for intracellular Mtb replication, it does not fully capture the unique features of pulmonary infection or potential drug-mutant interactions in the lungs. Therefore, we aimed to validate the identified synergistic CGIs using an aerosol infection model. We conducted aerosol infection of C57BL/6 mice with a selected mini-pool of 8 hypomorphs exhibiting synergistic CGIs across the four treatments (**Fig. 3A**) alongside three barcoded wild-type H37Rv strains. We adhered to the same experimental strategy as previously described, except for infection via the aerosol route (**Fig. 3B**). After two weeks of antibiotic treatment, we observed a significant reduction in lung (**Fig. 3C**) and spleen (**Fig. S3A**) bacterial burden across all treatment groups. We recovered, plated, and amplified barcode sequences from the surviving bacteria. Barcode counts were normalized to individual library size, and relative fitness was estimated by comparing barcode counts in groups that were DAS+4 depleted and drug-treated versus DAS+4 depleted in the absence of drug therapy. The relative fitness of each mutant in different treatment groups could then be compared to the pool of 3 independently barcoded clones of wild-type H37Rv (**Fig. 3D**). In this pulmonary infection model, we observed most of the synergistic interactions predicted in the larger screen. Knockdown of genes related to multiple functional pathways including translation (*aspS*, *rplE*, and *ettA*), cell wall biogenesis (*manB*), thiamine metabolism (*thiD*), and purine metabolism (*adk*) broadly increased the efficacy of the tested antibiotic regimens in the lungs. Notably, *qcrA* and *qcrB* had more modest effects than anticipated. Both genes are components of the same respiratory complex, and we speculate that their importance may vary between the lung and spleen. This is consistent with additional aerosol infection and treatment experiments in which the *qcrB* inhibitor Q203 (40) failed to show synergy with an INH/RIF combination regimen in the lungs (**Fig. S3B-C**). Regardless, we conclude that the CGI predictions from our screen are generally robust and reproducible during pulmonary infections, even though a fraction of the CGIs may display tissue-specific effects.

**Figure 3:**
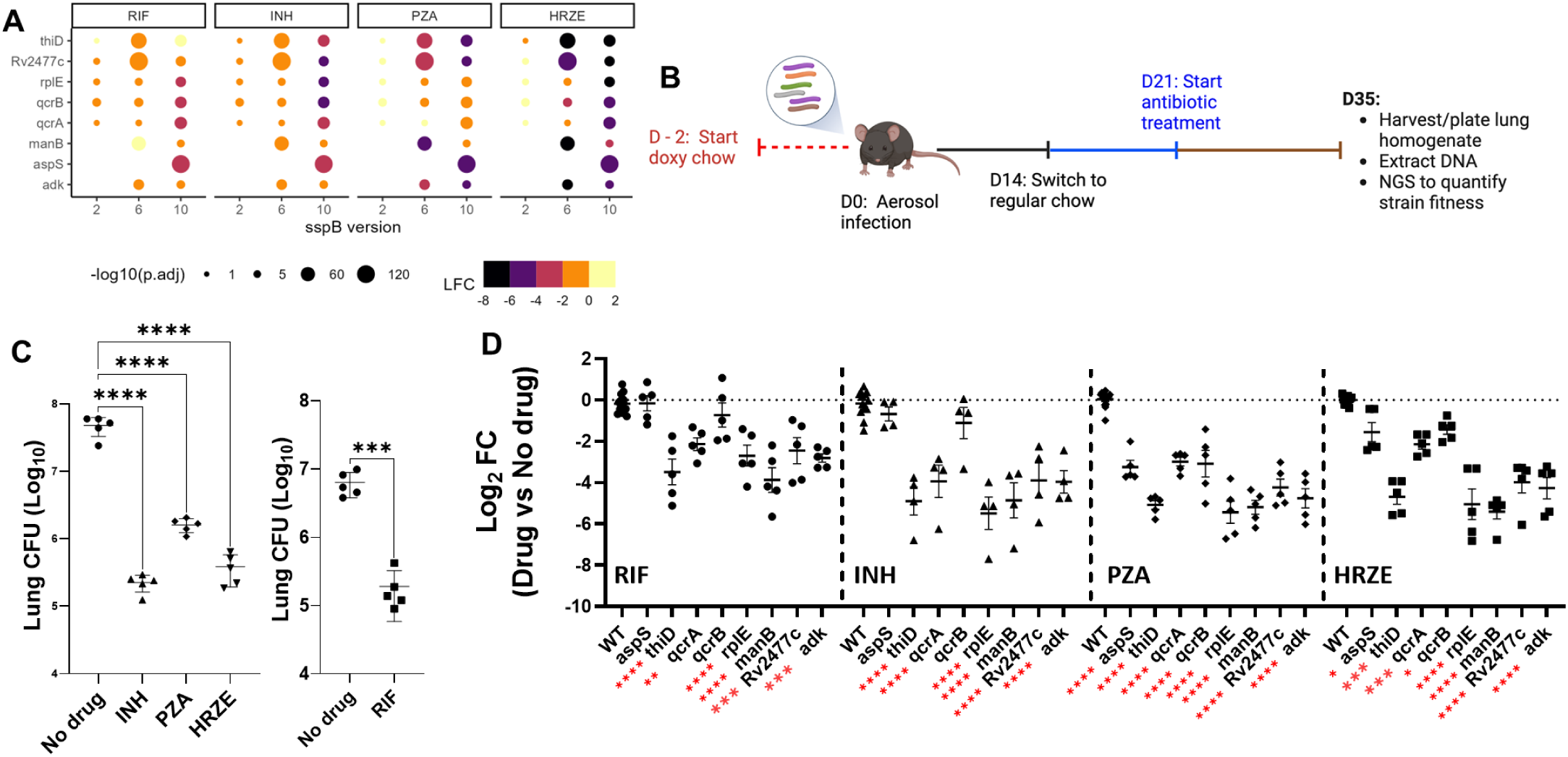
Top synergistic CGIs are reproducible during pulmonary mouse infection. **(A)** Bubble plot of the 8 hypomorph strains selected for aerosol infection mini pool. Plot size is scaled by the log_10_(FDR) values while the color range represents the log_2_ FC values. **(B)** Illustration of the mini-pool aerosol infection experiment for validation of top synergistic hits. **(C)** Right - Comparison of lung CFU in depleted-treated vs depleted-untreated mice groups (INH, PZA, and HRZE) (n = 5 biological replicates, mean ± s.e.m; One-way ANOVA with Dunnett’s multiple comparisons test; ****P < 0.0001). Left - Comparison of lung CFU in depleted-RIF vs depleted-untreated mice (n = 5 biological replicates, mean ± s.e.m; Two-tailed T-test; ***P < 0.001). **(D)** Comparison of relative abundance (log_2_ fold changes, treated vs non-treated) of different hypomorph strains vs wildtype H37Rv strains across different antibiotic treatment conditions. (n = 5 (hypomorphs) or n=15 (H37Rv) biological replicates, mean ± s.e.m). Results from Two-way ANOVA with Dunnett’s multiple comparisons test; *P < 0.05, **P < 0.01, ***P < 0.001, ****P < 0.0001.

### Diverse Mtb pathways alter antibiotic efficacy during infection

We next sought to identify bacterial pathways enriched in significant CGIs across different antibiotics in our dataset and to define novel mechanisms of antibiotic activity unique to the *in vivo* environment. First, we used hierarchical clustering of the LFC values to determine the relationship between mutants with significant synergistic CGIs in at least one condition (**Fig. 4A**). We observed patterns reflective of drug-specific and drug-independent interactions, with synergies that are apparent across multiple treatment conditions (drug-agnostic, cluster 2), those reflecting PZA or HRZE synergies (clusters 1 and 4), or those with more complex relationships between treatments (cluster 3).

**Figure 4:**
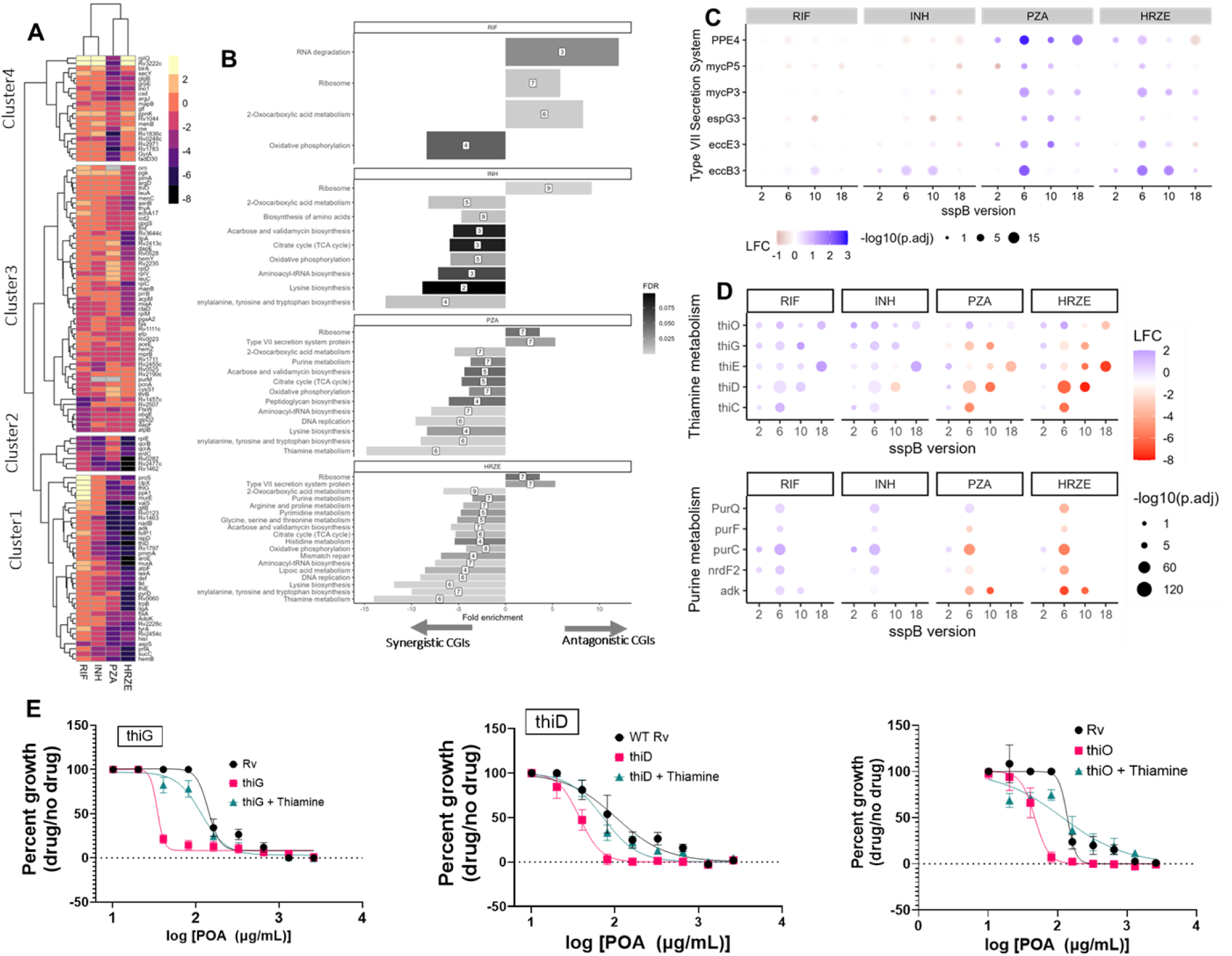
Diverse Mtb pathways alter antibiotic efficacy during infection. **(A)** Hierarchical clustering of log_2_FC values (treated vs non-treated) for mutants with log_2_FC < −1 and p.adj < 0.01 (synergistic CGIs) in at least one treatment condition (sspB6). **(B)** KEGG functional enrichment analysis of significant synergistic and antagonistic interactions across all treatment conditions. Numbers represent gene counts in each pathway **(C)** Bubble plots of log_2_ FC for ESX-3 Type VII secretion mutants across different conditions. **(D)** Bubble plots of log_2_ FC for mutants in thiamine and purine biosynthetic pathways. Plot sizes are scaled by the log_10_(FDR) values while the color range represents the log_2_ FC values. **(E)** Pyrazinoic acid (POA) dose-response curves across multiple thiamine knockdown mutants and wildtype H37Rv in the presence or absence of 100μM of thiamine. Data represent the average and standard error of the mean (SEM) growth (expressed as % growth relative to no drug growth controls in five biological replicates).

Next, we conducted functional enrichment analysis using a curated gene function annotation from Mtb Network Portal (41), BioCyc (42), and KEGG databases (43) to identify bacterial functions and pathways that alter *in vivo* antibiotic efficacy. We observed drug-specific CGIs that were consistent with the known mechanisms of action for these drugs (**Fig. 4B**). Consistent with rifampicin’s known ability to inhibit transcription, antagonistic GCIs with this drug were enriched for proteins associated with RNA degradation (**Fig. S3D**). For INH, synergistic CGIs related to central carbon metabolism (TCA cycle and 2-oxocarboxylic acid metabolism) could be related to this drug’s ability to inhibit long-chain fatty acid synthesis. We also noted pathways that consistently altered the efficacy of multiple drugs. Depletion in ribosomal proteins broadly reduced antibiotic efficacy in all treatment conditions (**Fig. 4B**). Similarly, most of the pathways enriched for isoniazid synergies, including amino acid and tRNA biosynthesis, were also enriched in PZA and HRZE conditions, highlighting the broad effects of these pathways on antibiotic efficacy. As both amino acid biosynthesis and tRNA synthetases are targets for first-in-class antibiotic drug discovery (44, 45), targeting these pathways could synergize broadly with current TB drugs.

### CGIs reveal mechanisms controlling PZA activity during infection

Despite its critical role in modern TB drug therapy, the mode(s) of action (MOA) of PZA during infection are still debated (46, 47). While we did not observe pathway enrichment for CoA biosynthesis, we did observe statistically significant synergy between PZA and its proposed enzymatic target, PanD (LFC -0.98, *P* = 0.03), along with another pathway member, PanB, (LFC -3.19, *P* < 0.001) (**Supplementary Table 1)**. In addition, we observed functional enrichment of several Mtb pathways that have not previously been implicated in PZA MOA (**Fig. 4B**). Disrupting genes in the ESX-3 type VII secretion system reduced *in vivo* PZA activity (**Fig. 4B**). Depletion of components of this system (*eccE3*, *espG*3, *eccB3*, *mycP3,* and PPE4) significantly reduced the efficacy of PZA (antagonistic interactions), and several of these mutants (eccB3, mycP3, and eccE3) showed a similar effect in the combination treatment condition (**Fig. 4C**). The conserved ESX-3 complex is involved in metal ion acquisition, particularly in mycobactin-mediated iron uptake (48). Intracellular iron has been shown to enhance the *in vitro* activity of PZA and Pyrazinoic acid (POA) against Mtb (49) by enhancing PncA enzyme activity and hydrolysis of PZA to POA (50). Thus, we hypothesize that disrupting ESX-3 components may reduce intracellular iron levels in Mtb, in the context of mouse infection, limiting PZA activation and decreasing *in vivo* antibiotic efficacy.

Additional functional enrichment analysis for overlapping synergistic interactions in PZA and HRZE conditions also identified multiple unique Mtb pathways. Mutants in the dedicated steps of thiamine metabolism, a pathway involved in the synthesis of the essential cofactor thiamine pyrophosphate (TPP), were enriched for synergistic CGIs in PZA and HRZE. Purine metabolism, which shares the first five enzymatic steps with thiamine metabolism, was also significantly enriched for synergistic PZA/HRZE interactions (**Fig. 4B**). Knockdowns in both thiamine and purine metabolic pathways consistently increased the *in vivo* efficacy of PZA and HRZE in the spleen and during pulmonary infection (**Fig. 3D**; **Fig. 4D**). To further validate the observed PZA synergy, we sought to characterize the effects of depleting thiamine genes on *in vitro* POA activity. Knocking down genes in the thiamine biosynthetic pathway increased POA sensitivity, with up to a 4-fold decrease in the IC_50_ values. Supplementing the media with 100µM of thiamine restored the *in vitro* POA sensitivity of these mutants to the wildtype levels (**Fig. 4E**). In sum, these data support PanD inhibition as an important mechanism of PZA activity during infection. In addition, as Mtb is unable to acquire sufficient thiamine and purine precursors from host tissues to support growth (51), both pathways could be targeted for the design of novel TB drugs that augment the effects of existing regimens.

## Discussion

Recent efforts to expand the TB drug pipeline have leveraged chemical-genetic interaction screens to identify promising targets for drug development. However, the influence of the host environment on these interactions remains largely unknown. Here, we leveraged an inducible knockdown library of essential Mtb genes to characterize determinants of antibiotic efficacy and uncover mechanisms of antibiotic action during infection. The findings from this work build on reports from previous studies leveraging drug-drug and drug-mutant interaction screens under *in vitro* culture conditions predictive of antibiotic efficacy in mouse models and human tuberculosis (13, 23). Our study furthers this effort by focusing on essential genes that are the most relevant targets for TB drug discovery and identifying CGIs directly in the infection environment.

We identified an *in vivo* CGI signature that is distinct from those previously identified using *in vitro* assays. Consistent with previous findings using Tn-Seq (32), no single *in vitro* condition or combination of *in vitro* conditions from this study served as a significant predictor of *in vivo* killing. We note that these previous studies differed from ours in multiple ways. In addition to the differences in environmental conditions, the *in vitro* studies detected changes in antibiotic MIC, whereas our studies focused on the rate of bacterial killing. While we are not able to attribute altered CGI profiles specifically to one of these experimental differences, it is clear that MIC changes in simple *in vitro* systems are a poor predictor of interactions that produce increased bacterial clearance during infection. More complex *in vitro* models or mathematical approaches that integrate many experimental conditions (23, 52) are likely necessary to adequately predict *in vivo* effects. The rich dataset of CGI from infected animals that we provide should serve as a useful resource to validate more predictive *in vitro* models.

Profiling drug-mutant interactions directly in the host environment can identify novel targets for combinatorial TB drug discovery. We identified multiple synergistic interactions, including mutants that result in drug-specific interactions and those with drug-agnostic interaction patterns. These interactions were independent of mutant growth rate, a common confounder of antibiotic activity (53, 54), suggesting that the observed interactions result from the individual effects of gene knockdown and antibiotic treatment. Most drug-specific synergies were also apparent in the HRZE multidrug treatment condition and could therefore be prioritized for the generation of efficacious combination regimens (24, 55). Drug-agnostic synergies are also promising targets for designing novel drugs that can be combined broadly with frontline TB drugs. We identified promiscuous synergies in 11 genes spanning multiple Mtb pathways. For example, disrupting the first committed step of *de novo* NAD biosynthesis, catalyzed by aspartate oxidase *nadB* (56), broadly synergized with all conditions. Mtb synthesizes nicotinamide adenine dinucleotide (NAD+) through both *de novo* and salvage pathways, with genes downstream of the crossroad for the two pathways, including *nadD* and *nadE*, identified as putative drug targets (56, 57). The broad synergistic effect that we describe for *nadB* suggests that *nadB* is also an attractive target for TB drug development. We also observed that disrupting components of protein translation, including tRNA synthetases, which are currently being explored as TB drug targets (58, 59), broadly synergized with different antibiotics during infection. Knockdown of *ettA*, an ABC-F protein involved in regulating the translation elongation cycle (60, 61), also increased the *in vivo* efficacy of different antibiotics. Naturally occurring mutations in *ettA* have been shown to constitutively upregulate *whiB7* stress response, causing low-level resistance to multiple antibiotics (62–64). Together, these observations suggest that alterations in the activity of *ettA* can have varied effects on antibiotic action.

PZA remains the cornerstone component of TB drug therapy and has been credited with shortening the treatment duration and improving treatment outcomes (65). However, the potency of this drug depends on conditions that are only fully reproduced in the infection environment, leaving the mechanisms underlying its clinical efficacy uncertain (46). Pyrazinoic acid (POA), the product of PZA hydrolysis by Mtb PncA enzyme, is known to inhibit coenzyme A (CoA) biosynthesis by targeting the aspartate decarboxylase, PanD (16, 47, 66). We observed that depleting either PanD or another enzyme in the pathway, PanB, increased *in vivo* PZA sensitivity, supporting the importance of this mechanism of action. We also observed that disrupting multiple other bacterial pathways influenced PZA efficacy. For instance, mutants of the ESX-3 type VII secretion system, a specialized secretion system implicated in mycobactin-mediated iron uptake (48, 67), decreased the *in vivo* PZA efficacy. Fe^2+^ plays a catalytic role in pncA enzymatic activity, enhancing the hydrolysis of PZA to POA and increasing the *in vitro* activity of PZA/POA against Mtb (49, 50). Even though the ESX-3 system is relatively conserved in Mtb clinical isolates (68), *pncA* mutations associated with PZA resistance have been mapped to key interaction sites, including iron coordination residues (amino acids 49, 51, 57, and 71) (69, 70). Thus, reduced intracellular Fe^2+^ levels could impair the catalytic activity of *pncA* and reduce PZA efficacy during infection.

We also observed PZA-specific synergies with mutants in thiamine and purine biosynthetic pathways, suggesting a potential link between these pathways and PZA activity. Recent studies have demonstrated that Mtb requires *de novo* purine and thiamine biosynthesis for survival during infection (51). Thiamine pyrophosphate (TPP), the active form of thiamine, is a crucial cofactor for enzymes in various metabolic pathways (71, 72), making its biosynthetic pathway a promising target for antibiotic development (73–76). Studies in *Salmonella enterica* have described crosstalk between CoA, and thiamine, and purine biosynthesis that could explain the observed CGI between these pathways and PZA. Disrupting pantothenate biosynthesis in *Salmonella* not only reduces intracellular CoA levels but also causes conditional thiamine auxotrophy when purine biosynthesis is compromised (77–79). Though thiamine biosynthesis is not as well characterized in Mtb, these studies in *Salmonella* suggest that conditions that decrease intracellular CoA levels, including PZA/POA treatment, increase bacterial dependence on the interconnected thiamine and purine metabolic pathways. Thus, both thiamine and purine biosynthesis may be attractive targets for developing synergistic TB drug combinations.

This work is not without caveats. First, although the short-term treatment we employed in the mouse model reflects the early bactericidal activity (EBA) approach used in clinical trials, it is not necessarily predictive of sterilizing activity (80). Second, while our hypomorph system can recapitulate partial chemical inhibition effects, genetic knockdown is not the same as pharmacological modulation, which can have complex effects beyond simple enzyme inhibition. Lastly, even though the CGIs we identified in splenic infection largely reflect interactions during pulmonary infection, they do not perfectly model tuberculosis infection at this site, as we observed with *qcrAB* mutants. Despite these caveats, our broad and unbiased characterization of *in vivo* drug-mutant interactions elucidated mechanisms of antibiotic activity during infection that will facilitate the development of more predictive *in vitro* models and the prioritization of more promising drug candidates.

## Materials and Methods

### Bacterial cultures

All strains were generated from the Mtb H37Rv parent strain, and all experimental procedures followed CDC-NIH biosafety guidelines. Cells were grown in standard DIFCO Middlebrook 7H9 broth with 10% Oleic acid-albumin-dextrose-catalase (OADC), 0.2% glycerol and 0.05% Tween-80 and platted in 7H10 agar with 10% OADC, 0.5% glycerol and appropriate antibiotics. ATc-inducible hypomorph strains were grown in appropriate antibiotics and supplemented with 500ng/mL anhydrotetracycline (ATc). All hypomorph strains included in the mouse intravenous and aerosol infection pool were cultured to the mid-log phase followed by washing and back-dilution to an OD_600_ of 0.7 before pooling at 100 colony-forming units (CFUs) per strain per depletion level on the day of mouse infection.

### Antibiotics quantification in drinking water and plasma pharmacokinetics

Healthy C57BL/6J mice (four mice per group) were administered rifampicin (RIF, Millipore-Sigma R3501), isoniazid (INH, Millipore-Sigma 54-85-3), pyrazinamide (PZA, Millipore-Sigma 98-96-4), and ethambutol (EMB, Millipore-Sigma 1070-11-7) in drinking water alone (0.5g/L INH, 0.1g/L RIF, 0.6g/L EMB or 1.5g/L PZA) or in HRZE combination (0.1g/L INH, 0.1g/L RIF, 1.2g/L EMB or 1.5g/L PZA). On the third day, serial blood samples were collected first thing in the morning (∼7:30AM) for 5 consecutive days for plasma pharmacokinetics and stored at -80℃. To quantify antibiotic stability in drinking water alone or in the HRZE combination, daily duplicate water samples were collected for 7 days. Antibiotic concentration in drinking water and plasma pharmacokinetic analysis was done as previously described (13) using high-performance liquid chromatography-tandem mass spectrometry (HPLC-MS/MS). INH, PZA, and EMB alone and in the HRZE combination remained stable in drinking water for 7 days. However, RIF was slightly unstable when administered alone and displayed increased instability in the HRZE combination. Thus, RIF and HRZE antibiotic water were replaced every other day, while INH and PZA water were replaced weekly.

### In vivo chemical-genetic interaction screening

Male C57BL/6J mice were purchased from the Jackson Laboratory (Bar Harbor, ME, USA). All experiments and housing were conducted in conformity with the Department of Animal Medicine of the University of Massachusetts Chan Medical School and Institutional Animal Care and Use Committee (IACUC) guidelines. All mice were started on 2000 ppm doxycycline rodent chow (Research Diets, New Brunswick, NJ, USA) two days before intravenous injection (IV) with a pool of hypomorph mutants at 100 CFU per strain (∼50,000 CFU). The no-depletion, no-treatment control groups were maintained on a doxycycline diet for the duration of the study. At 14 days post-infection (14dpi), all the other groups were switched to the regular diet, with daily cage changes for one week to prevent mice from consuming residual doxycycline via coprophagy. At 21dpi, antibiotics were administered via drinking water to the indicated groups at the following optimized concentrations: 0.5g/liter INH, 0.1g/liter RIF, 1.5g/liter PZA and HRZE combination cocktail (0.1g/L INH + 0.1g/liter RIF + 1.2g/liter EMB + 1.5g/liter PZA). At the end of each time point, all mice were euthanized, and surviving bacteria were recovered from both the spleen and lungs followed by plating on 7H10 agar plates supplemented with 500ng/mL ATC and appropriate antibiotics. Approximately 10^6 CFU/mouse were plated for library recovery and subsequent sequencing to estimate strain abundance.

### Aerosol infection validation experiment

To validate hits from the screen, aerosol infection of male C57BL/6J mice with a pool of 11 strains (inclusive of three barcoded wild-type strains) at ∼1100 CFU/mouse, was conducted. We followed the same protocol, timelines, and antibiotic concentrations as described in the previous section.

### DNA extraction, library preparation, Illumina sequencing, and analysis

Genomic DNA was extracted from recovered bacterial pellets following BSL3 guidelines using the phenol-chloroform-isoamyl alcohol method. Briefly, Mtb pellets scraped from agar plates were resuspended in a mixture of 10mL of 0.01M TE buffer (0.01M Tris-HCL, 1mM EDTA, pH9) and 10mL of phenol-chloroform-isoamyl alcohol (25:24:1). The cell suspensions were centrifuged at 4000 rpm and the pellet was recovered by removing the aqueous and organic phases. The dried bacterial mass was resuspended in 5mL of 0.1M TE buffer (0.1M Tris-HCL, 1mM EDTA, pH9), vortexed, and 100 uL of 5mg/mL lysozyme added (Millipore-Sigma, 9066-59-5) before incubation at 37°C for 14-16 hours. Next, 500uL of 10% SDS (Millipore-Sigma, 151-21-3) and 28uL of 20mg/mL proteinase K (Invitrogen, 25530049) were added and incubated for 3-4 hours at 50-56°C. Samples were then treated with 6mL of 1:1 phenol: chloroform for 30 minutes at room temperature before being safely removed from the BSL3 facility. All samples were then centrifuged at 4000 rpm before transferring the aqueous layer to phase-locked tubes. Each sample was mixed with 5mL of 1:1 phenol:chloroform, centrifuged, and aqueous layer recovered in new tubes. DNA was precipitated from the clear aqueous layer (300-800 µl) using 1/10 volume of 3M sodium acetate and 1 volume of isopropanol, washed twice with 80% ethanol, dried, and resuspended in 0.1M TE buffer.

Extracted genomic DNA was quantified using NanoDrop 2000/200c (ThermoFisher Scientific, ND-2000). Next, barcode junctions for each hypomorph strain were amplified from 40ng genomic DNA with 22 cycles using Q5 High Fidelity 2X master mix (NEB M0492L). Each PCR reaction was done in 40μL containing 4μL genomic DNA, 6μL of 10μM primer pairs (IDT), 20μL Q5 High Fidelity 2X master mix, and 4µL of DNase-free molecular grade water. Primers were designed to include Truseq Dual indexing and custom stagger sequences to improve base diversity for sequencing. Amplified libraries were purified using AmpureXP beads and quantified by Qubit 1x dsDNA HS Assay Kit (Invitrogen, Q33231). A final 4nM equimolar pool of each library was made, followed by qPCR to confirm library concentration using the Kappa Library Quantification Kit (Roche, KK4824). 5-10% phiX spike-in (NextSeq PhiX Control Kit, Illumina) was added to the library to further support sequencing diversity and samples were run on the Illumina NextSeq 1000/2000 platform (single-end, dual-indexed, 1×100 cycles). Barcode counts extracted from high-quality sequencing reads using an in-house bash script were analyzed using DEBRA (39) to identify differentially represented barcodes (DRBs) in different experimental mice groups, reporting the log_2_ fold changes and FDR-adjusted *P* values. For the aerosol validation mini-pool experiment, the relative barcode abundance for each strain was calculated as the mean ratio of barcode counts of each strain relative to the three wild-type strains in the pool.

### Differential barcode representation analysis

To define synergistic, antagonistic, and non-significant chemical-genetic interactions, extracted barcode counts in depleted-treated vs depleted-no treatment were compared using DEBRA (39). DEBRA implements a modified version of DESeq2 analysis (81) optimized for identifying differentially represented barcodes. We implemented Wald statistical tests with significant CGI cutoff at log2FC |1| and false discovery rate (FDR) adjusted *P* value of 0.01. A barcode filtering threshold was designed to exclude low-abundance hypomorphs lost from the pool due to infection bottlenecks. Mutants with consistently low barcode counts (less than 10 counts) in both infection controls (1-dpi) and no-treatment controls (35-dpi) were discarded from further analysis.

### In vitro antibiotic susceptibility

For *in vitro* MIC testing, all hypomorphs and wild-type strains were inoculated in 7H9 medium with OADC, 0.2% glycerol, and 0.05% Tween-80 at OD_600_ of 0.05 with two-fold serial dilutions of POA (pyrazinecarboxylic acid; Millipore-Sigma, 98-97-5). Where indicated, 100μM final concentration of thiamine hydrochloride (Millipore-Sigma, 67-03-8) was added. All conditions and strains were assessed in quadruplets and growth was monitored by OD_600_ measurements. Percent growth was estimated as OD_600_ at different antibiotic concentrations relative to the growth (no drug) control.

### Q203 mouse experiments

To determine the efficacy of Q203 alone or with isoniazid and rifampicin combination, we conducted an aerosol mouse infection and treatment experiment. Briefly, mice were infected with wildtype *M. tuberculosis* H37Rv at ∼100 CFU per lung. Treatment started 4 weeks after infection. Q203 (MedChem, 1334719-95-7) was formulated in 20% TPGS (D-α tocopheryl polyethylene glycol 1000 succinate; MedChem, 9002-96-4) and administered daily via oral gavage at a dose of 2mg per kg body weight. INH/RIF treatment was administered orally at the previously used doses. All non-Q203 treated mice and controls also received daily oral gavage of the vehicle. Bacterial load in the lungs of all mice was determined after 2 weeks of treatment by CFU enumeration.

## Acknowledgments

We are grateful to past and present members of the Sassetti lab for their valuable discussions, critiques, and assistance throughout this work. This work was supported by grants from the National Institutes of Health (AI162598 and AI095208).

## Data availability

All data generated in this study are included within the manuscript and in Supplemental Information. Raw barcode sequencing data will be available on the NCBI Short Read Archive (SRA) (Project and accession number will be provided before publication).

## Code availability

The custom code used in the analysis of this study is available at https://github.com/Owuorgpo/In-vivo-CGI

## Bibliography

1. World Health Organization, Global tuberculosis report 2023 (World Health Organization, 2023).

2. W. Fox, D. A. Mitchison, Short-course chemotherapy for pulmonary tuberculosis. Am. Rev. Respir. Dis. 111, 325–353 (1975).

3. R. Colangeli, et al., Bacterial Factors That Predict Relapse after Tuberculosis Therapy. N. Engl. J. Med. 379, 823–833 (2018).

4. F. Conradie, et al., Treatment of Highly Drug-Resistant Pulmonary Tuberculosis. N. Engl. J. Med. 382, 893–902 (2020).

5. B.-T. Nyang’wa, et al., A 24-Week, All-Oral Regimen for Rifampin-Resistant Tuberculosis. N. Engl. J. Med. 387, 2331–2343 (2022).

6. M. D. J Libardo, H. I. Boshoff, C. E. Barry 3rd, The present state of the tuberculosis drug development pipeline. Curr. Opin. Pharmacol. 42, 81–94 (2018).

7. N. Ismail, S. V. Omar, N. A. Ismail, R. P. H. Peters, Collated data of mutation frequencies and associated genetic variants of bedaquiline, clofazimine and linezolid resistance in Mycobacterium tuberculosis. Data Brief 20, 1975–1983 (2018).

8. S. Kadura, et al., Systematic review of mutations associated with resistance to the new and repurposed Mycobacterium tuberculosis drugs bedaquiline, clofazimine, linezolid, delamanid and pretomanid. J. Antimicrob. Chemother. 75, 2031–2043 (2020).

9. N. Ismail, et al., Genetic variants and their association with phenotypic resistance to bedaquiline in Mycobacterium tuberculosis: a systematic review and individual isolate data analysis. Lancet Microbe 2, e604–e616 (2021).

10. S.-H. Baek, A. H. Li, C. M. Sassetti, Metabolic regulation of mycobacterial growth and antibiotic sensitivity. PLoS Biol. 9, e1001065 (2011).

11. Y. Liu, et al., Immune activation of the host cell induces drug tolerance in Mycobacterium tuberculosis both in vitro and in vivo. J. Exp. Med. 213, 809–825 (2016).

12. J. E. Beam, S. E. Rowe, B. P. Conlon, Shooting yourself in the foot: How immune cells induce antibiotic tolerance in microbial pathogens. PLoS Pathog. 17, e1009660 (2021).

13. E.-I. Koh, et al., Chemical-genetic interaction mapping links carbon metabolism and cell wall structure to tuberculosis drug efficacy. Proc. Natl. Acad. Sci. U. S. A. 119, e2201632119 (2022).

14. Short-course chemotherapy in pulmonary tuberculosis. A controlled trial by the British Thoracic and Tuberculosis Association. Lancet 2, 1102–1104 (1976).

15. E. A. Lamont, A. D. Baughn, Impact of the host environment on the antitubercular action of pyrazinamide. EBioMedicine 49, 374–380 (2019).

16. P. Gopal, et al., Pyrazinoic Acid Inhibits Mycobacterial Coenzyme A Biosynthesis by Binding to Aspartate Decarboxylase PanD. ACS Infect Dis 3, 807–819 (2017).

17. C. M. Sassetti, E. J. Rubin, Genetic requirements for mycobacterial survival during infection. Proc. Natl. Acad. Sci. U. S. A. 100, 12989–12994 (2003).

18. C. M. Sassetti, D. H. Boyd, E. J. Rubin, Genes required for mycobacterial growth defined by high density mutagenesis. Mol. Microbiol. 48, 77–84 (2003).

19. N. Lyons, W. Wu, Y. Jin, I. L. Lamont, D. Pletzer, Using host-mimicking conditions and a murine cutaneous abscess model to identify synergistic antibiotic combinations effective against. Front. Cell. Infect. Microbiol. 14, 1352339 (2024).

20. V. Eldholm, F. Balloux, Antimicrobial Resistance in Mycobacterium tuberculosis: The Odd One Out. Trends Microbiol. 24, 637–648 (2016).

21. C. A. Kerantzas, W. R. Jacobs Jr, Origins of Combination Therapy for Tuberculosis: Lessons for Future Antimicrobial Development and Application. MBio 8 (2017).

22. M. Zhu, M. W. Tse, J. Weller, J. Chen, P. C. Blainey, The future of antibiotics begins with discovering new combinations. Ann. N. Y. Acad. Sci. 1496, 82–96 (2021).

23. J. Larkins-Ford, Y. N. Degefu, N. Van, A. Sokolov, B. B. Aldridge, Design principles to assemble drug combinations for effective tuberculosis therapy using interpretable pairwise drug response measurements. Cell Rep Med 3, 100737 (2022).

24. M. Tyers, G. D. Wright, Drug combinations: a strategy to extend the life of antibiotics in the 21st century. Nat. Rev. Microbiol. 17, 141–155 (2019).

25. J. Pieters, J. D. McKinney, Pathogenesis of Mycobacterium tuberculosis and its Interaction with the Host Organism (Springer Science & Business Media, 2013).

26. S. Ma, et al., Transcriptomic Signatures Predict Regulators of Drug Synergy and Clinical Regimen Efficacy against Tuberculosis. MBio 10 (2019).

27. I. Katzir, M. Cokol, B. B. Aldridge, U. Alon, Prediction of ultra-high-order antibiotic combinations based on pairwise interactions. PLoS Comput. Biol. 15, e1006774 (2019).

28. J. M. Rock, et al., Programmable transcriptional repression in mycobacteria using an orthogonal CRISPR interference platform. Nat Microbiol 2, 16274 (2017).

29. J.-H. Kim, et al., Protein inactivation in mycobacteria by controlled proteolysis and its application to deplete the beta subunit of RNA polymerase. Nucleic Acids Res. 39, 2210–2220 (2011).

30. J. E. Long, et al., Identifying essential genes in Mycobacterium tuberculosis by global phenotypic profiling. Methods Mol. Biol. 1279, 79–95 (2015).

31. W. Xu, et al., Chemical Genetic Interaction Profiling Reveals Determinants of Intrinsic Antibiotic Resistance in Mycobacterium tuberculosis. Antimicrob. Agents Chemother. 61 (2017).

32. M. M. Bellerose, et al., Distinct Bacterial Pathways Influence the Efficacy of Antibiotics against Mycobacterium tuberculosis. mSystems 5 (2020).

33. E. O. Johnson, et al., Large-scale chemical-genetics yields new M. tuberculosis inhibitor classes. Nature 571, 72–78 (2019).

34. B. B. Mishra, et al., Nitric oxide prevents a pathogen-permissive granulocytic inflammation during tuberculosis. Nat Microbiol 2, 17072 (2017).

35. C. M. Smith, et al., Host-pathogen genetic interactions underlie tuberculosis susceptibility in genetically diverse mice. Elife 11 (2022).

36. Y. J. Zhang, et al., Tryptophan biosynthesis protects mycobacteria from CD4 T-cell-mediated killing. Cell 155, 1296–1308 (2013).

37. A. Tostmann, et al., Pharmacokinetics of first-line tuberculosis drugs in Tanzanian patients. Antimicrob. Agents Chemother. 57, 3208–3213 (2013).

38. S. Abel, P. Abel zur Wiesch, B. M. Davis, M. K. Waldor, Analysis of Bottlenecks in Experimental Models of Infection. PLoS Pathog. 11, e1004823 (2015).

39. Y. Akimov, D. Bulanova, S. Timonen, K. Wennerberg, T. Aittokallio, Improved detection of differentially represented DNA barcodes for high-throughput clonal phenomics. Mol. Syst. Biol. 16, e9195 (2020).

40. K. Pethe, et al., Discovery of Q203, a potent clinical candidate for the treatment of tuberculosis. Nat. Med. 19, 1157–1160 (2013).

41. S. Ma, et al., Integrated Modeling of Gene Regulatory and Metabolic Networks in Mycobacterium tuberculosis. PLoS Comput. Biol. 11, e1004543 (2015).

42. P. D. Karp, et al., The BioCyc collection of microbial genomes and metabolic pathways. Brief. Bioinform. 20, 1085–1093 (2019).

43. M. Kanehisa, M. Furumichi, M. Tanabe, Y. Sato, K. Morishima, KEGG: new perspectives on genomes, pathways, diseases and drugs. Nucleic Acids Res. 45, D353–D361 (2017).

44. E. J. Hasenoehrl, et al., Derailing the aspartate pathway of Mycobacterium tuberculosis to eradicate persistent infection. Nat. Commun. 10, 4215 (2019).

45. A. H. Diacon, et al., A first-in-class leucyl-tRNA synthetase inhibitor, ganfeborole, for rifampicin-susceptible tuberculosis: a phase 2a open-label, randomized trial. Nat. Med. 30, 896–904 (2024).

46. E. A. Lamont, N. A. Dillon, A. D. Baughn, The Bewildering Antitubercular Action of Pyrazinamide. Microbiol. Mol. Biol. Rev. 84 (2020).

47. Q. Sun, et al., The molecular basis of pyrazinamide activity on Mycobacterium tuberculosis PanD. Nat. Commun. 11, 339 (2020).

48. M. S. Siegrist, et al., Mycobacterial Esx-3 is required for mycobactin-mediated iron acquisition. Proc. Natl. Acad. Sci. U. S. A. 106, 18792–18797 (2009).

49. A. Somoskovi, M. M. Wade, Z. Sun, Y. Zhang, Iron enhances the antituberculous activity of pyrazinamide. J. Antimicrob. Chemother. 53, 192–196 (2004).

50. H. Zhang, et al., Characterization of Mycobacterium tuberculosis nicotinamidase/pyrazinamidase. FEBS J. 275, 753–762 (2008).

51. A. M. Block, P. C. Wiegert, S. B. Namugenyi, A. D. Tischler, Transposon sequencing reveals metabolic pathways essential for Mycobacterium tuberculosis infection. PLoS Pathog. 20, e1011663 (2024).

52. J. Larkins-Ford, et al., Systematic measurement of combination-drug landscapes to predict in vivo treatment outcomes for tuberculosis. Cell Syst 12, 1046–1063.e7 (2021).

53. R. H. Eng, F. T. Padberg, S. M. Smith, E. N. Tan, C. E. Cherubin, Bactericidal effects of antibiotics on slowly growing and nongrowing bacteria. Antimicrob. Agents Chemother. 35, 1824–1828 (1991).

54. J. Sarathy, V. Dartois, T. Dick, M. Gengenbacher, Reduced drug uptake in phenotypically resistant nutrient-starved nonreplicating Mycobacterium tuberculosis. Antimicrob. Agents Chemother. 57, 1648–1653 (2013).

55. V. A. Dartois, E. J. Rubin, Anti-tuberculosis treatment strategies and drug development: challenges and priorities. Nat. Rev. Microbiol. 20, 685–701 (2022).

56. C. Vilchèze, B. Weinrick, K.-W. Wong, B. Chen, W. R. Jacobs Jr, NAD+ auxotrophy is bacteriocidal for the tubercle bacilli. Mol. Microbiol. 76, 365–377 (2010).

57. S. Y. Gerdes, et al., From genetic footprinting to antimicrobial drug targets: examples in cofactor biosynthetic pathways. J. Bacteriol. 184, 4555–4572 (2002).

58. T. R. Ioerger, et al., Identification of new drug targets and resistance mechanisms in Mycobacterium tuberculosis. PLoS One 8, e75245 (2013).

59. R. Soto, et al., Identification and characterization of aspartyl-tRNA synthetase inhibitors against Mycobacterium tuberculosis by an integrated whole-cell target-based approach. Sci. Rep. 8, 12664 (2018).

60. G. Boël, et al., The ABC-F protein EttA gates ribosome entry into the translation elongation cycle. Nat. Struct. Mol. Biol. 21, 143–151 (2014).

61. Z. Cui, et al., Interplay between an ATP-binding cassette F protein and the ribosome from Mycobacterium tuberculosis. Nat. Commun. 13, 432 (2022).

62. K. Faksri, et al., Whole-Genome Sequencing Analysis of Serially Isolated Multi-Drug and Extensively Drug Resistant Mycobacterium tuberculosis from Thai Patients. PLoS One 11, e0160992 (2016).

63. S. Li, et al., CRISPRi chemical genetics and comparative genomics identify genes mediating drug potency in Mycobacterium tuberculosis. Nat Microbiol 7, 766–779 (2022).

64. L. K. R. Sharkey, T. A. Edwards, A. J. O’Neill, ABC-F Proteins Mediate Antibiotic Resistance through Ribosomal Protection. MBio 7, e01975 (2016).

65. W. Fox, D. A. Mitchison, Short-course chemotherapy for pulmonary tuberculosis. Am. Rev. Respir. Dis. 111, 845–8 CONTD (1975).

66. W. Shi, et al., Aspartate decarboxylase (PanD) as a new target of pyrazinamide in Mycobacterium tuberculosis. Emerg. Microbes Infect. 3, e58 (2014).

67. M. S. Siegrist, et al., Mycobacterial Esx-3 requires multiple components for iron acquisition. MBio 5, e01073–14 (2014).

68. Á. Chiner-Oms, M. G. López, M. Moreno-Molina, V. Furió, I. Comas, Gene evolutionary trajectories in reveal temporal signs of selection. Proc. Natl. Acad. Sci. U. S. A. 119, e2113600119 (2022).

69. S. Petrella, et al., Crystal structure of the pyrazinamidase of Mycobacterium tuberculosis: insights into natural and acquired resistance to pyrazinamide. PLoS One 6, e15785 (2011).

70. M. Karmakar, et al., Analysis of a Novel pncA Mutation for Susceptibility to Pyrazinamide Therapy. Am. J. Respir. Crit. Care Med. 198, 541–544 (2018).

71. C. K. Singleton, P. R. Martin, Molecular mechanisms of thiamine utilization. Curr. Mol. Med. 1, 197–207 (2001).

72. R. A. W. Frank, F. J. Leeper, B. F. Luisi, Structure, mechanism and catalytic duality of thiamine-dependent enzymes. Cell. Mol. Life Sci. 64, 892–905 (2007).

73. H. J. Kim, et al., The ThiL enzyme is a valid antibacterial target essential for both thiamine biosynthesis and salvage pathways in. J. Biol. Chem. 295, 10081–10091 (2020).

74. X. W. A. Chan, et al., Chemical and genetic validation of thiamine utilization as an antimalarial drug target. Nat. Commun. 4, 2060 (2013).

75. H. J. Kim, et al., Pharmacological perturbation of thiamine metabolism sensitizes Pseudomonas aeruginosa to multiple antibacterial agents. Cell Chem Biol 29, 1317–1324.e5 (2022).

76. L. A. Carfrae, E. D. Brown, Nutrient stress is a target for new antibiotics. Trends Microbiol. 31, 571–585 (2023).

77. D. M. Downs, L. Petersen, apbA, a new genetic locus involved in thiamine biosynthesis in Salmonella typhimurium. J. Bacteriol. 176, 4858–4864 (1994).

78. M. E. Frodyma, D. Downs, ApbA, the ketopantoate reductase enzyme of Salmonella typhimurium is required for the synthesis of thiamine via the alternative pyrimidine biosynthetic pathway. J. Biol. Chem. 273, 5572–5576 (1998).

79. D. C. Ernst, A. J. Borchert, D. M. Downs, Perturbation of the metabolic network in Salmonella enterica reveals cross-talk between coenzyme A and thiamine pathways. PLoS One 13, e0197703 (2018).

80. A. Jindani, V. R. Aber, E. A. Edwards, D. A. Mitchison, The early bactericidal activity of drugs in patients with pulmonary tuberculosis. Am. Rev. Respir. Dis. 121, 939–949 (1980).

81. M. I. Love, W. Huber, S. Anders, Moderated estimation of fold change and dispersion for RNA-seq data with DESeq2. Genome Biol. 15, 550 (2014).

