## Supplemental figures for "Chemical genetic interactions elucidate pathways controlling tuberculosis antibiotic efficacy during infection"

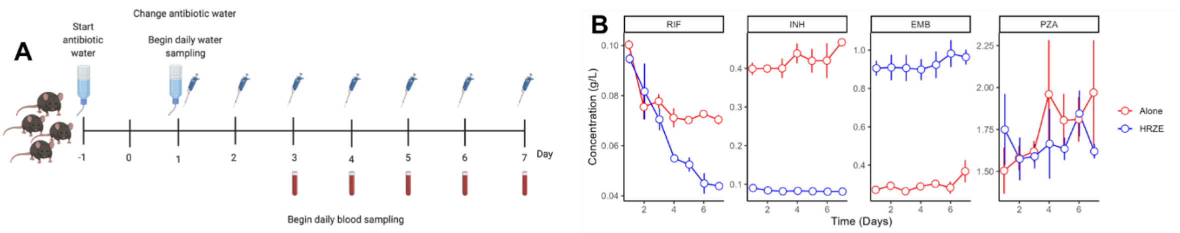


**C**

| | Concentration of antibiotic in water (mg/ml) | | 24-hour projected AUC (h*ug/ml) from mouse plasma | | Target human AUC (hr*ug/ml) range |
|---|---|---|---|---|---|
| | Alone | In HRZE | Alone | In HRZE | |
| INH | 0.5 | 0.1 | 52 ± 15 | 6 ± 2 | 4–23 |
| RIF | 0.1 | 0.1 | 82 ± 42 | 11 ± 7 | 27-68 |
| PZA | 1.5 | 1.5 | 320 ± 55 | 278 ± 66 | 209-610 |
| EMB | 0.6 | 1.2 | 20 ± 9 | 35 ± 10 | 13-32 |

**Supp. Figure 1: (A)** Schematic illustrating serial sampling and measurement of antibiotic stability (alone or in HRZE) from drinking water **(B)** Stability of different antibiotics in drinking water administered alone or in HRZE (n = 3, mean ± s.e.m). **(C)** Plasma concentrations of each antibiotic are represented by the average and standard deviation of the 24-hour *ug/ml concentration from 4 mice in each group. Target human area under curve (AUC) values are based on previously published values (38).

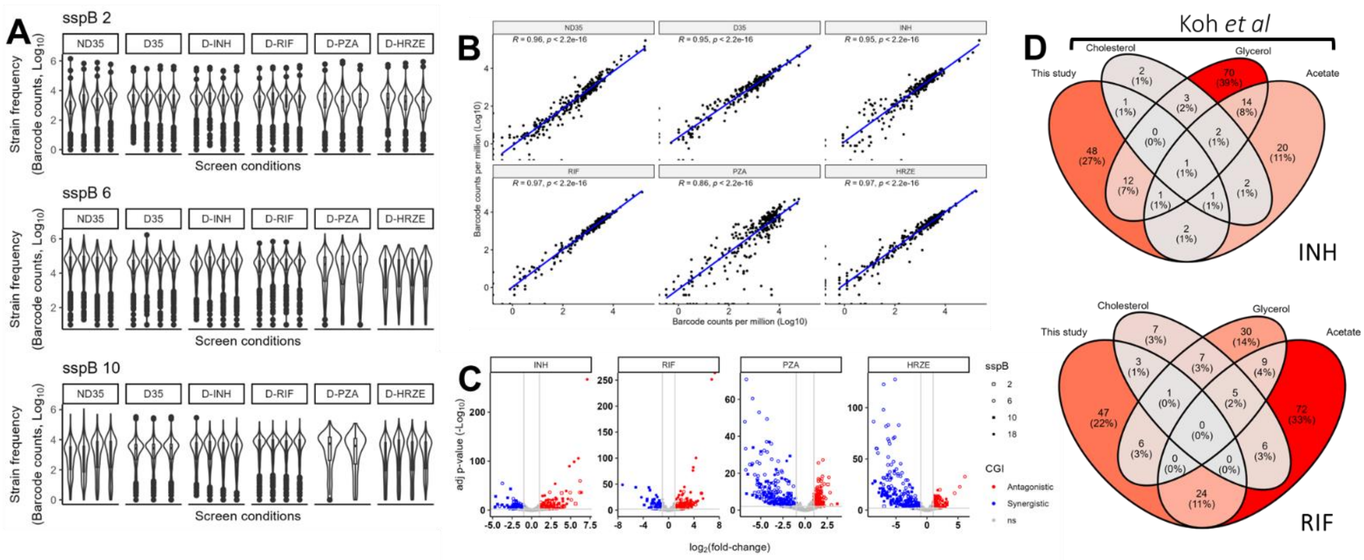


**Supp. Figure 2: (A)** Violin plots depicting the number of times each mutant was detected in each experimental condition across different depletion levels. **(B)** Spearman correlation of normalized barcode counts (reads per million, RPM) in biological replicates from different experimental conditions. Nondepleted (ND35), Depleted, no treatment (D35), and Depleted and treated (D-INH, D-RIF, D-PZA, and D-HRZE). **(C)** Volcano plot of interactions across all treatment groups and depletion levels during infection. The sspB represents the different depletion levels; ns (not significant) **(D)** Overlap of antagonistic interactions across different *in vitro* culture conditions (13) and during infection in INH and RIF conditions.

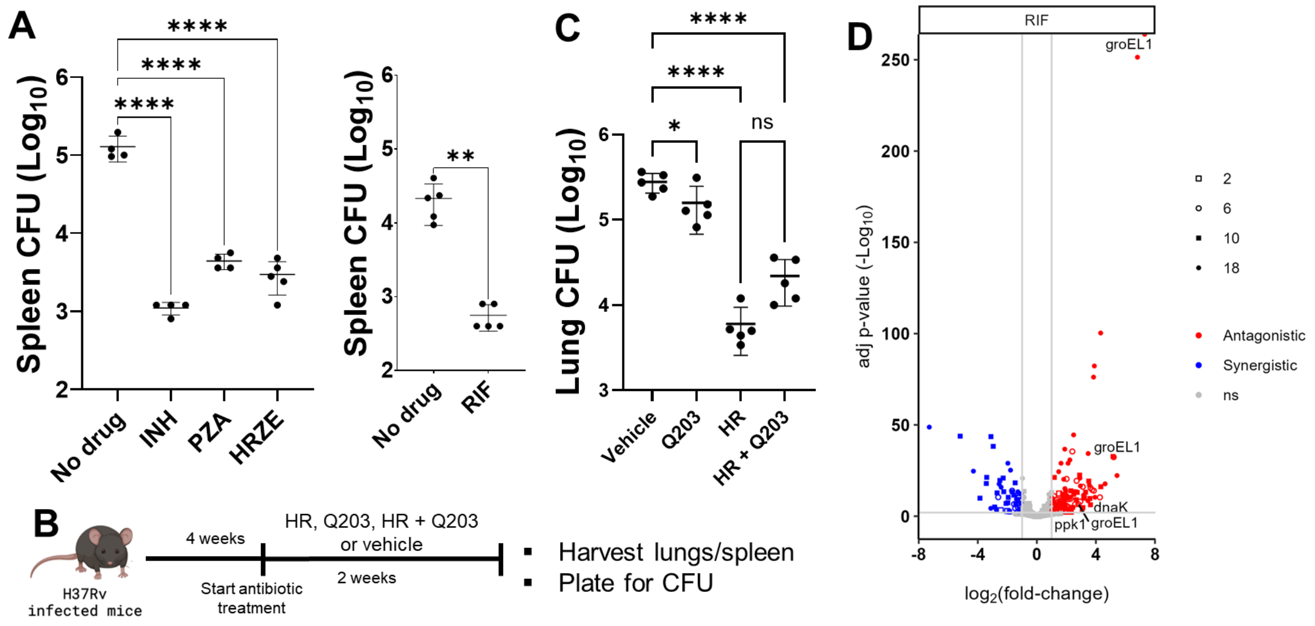


**Supp. Figure 3**: **(A)** Spleen CFU in depleted-treated vs depleted-untreated mice groups (n = 5 biological replicates, mean ± s.e.m; One-way ANOVA with Dunnett's multiple comparisons test or Two-tailed T-test; ****$P < 0.0001$, **$P < 0.01$). **(B)** Experimental design of mouse infection and treatment with HR, Q203 alone, HR + Q203, or vehicle (see methods). **(C)** Comparison of lung CFU in vehicle vs mice treated with Q203, HR combination, or HR + Q203 combination. HR (Isoniazid and rifampicin), Q203 (Telacebec; imidazopyridine amide), HR + Q203 (INH/RIF/Q203 combination). **(D)** Volcano plot of rifampicin interactions, highlighting antagonistic interactions in RNA degradation proteins. Points are shaped by depletion levels and colored by type of interaction.
